# Functional integrity of visual coding following advanced photoreceptor degeneration

**DOI:** 10.1101/2022.07.27.501697

**Authors:** Jessica Rodgers, Steven Hughes, Moritz Lindner, Annette E Allen, Aghileh Ebrahimi, Riccardo Storchi, Stuart N Peirson, Robert J Lucas, Mark W Hankins

## Abstract

Photoreceptor degeneration sufficient to produce severe visual loss often spares the inner retina. This raises the hope that treatments using optogenetics or electrical stimulation, which generate a replacement light input signal in surviving neurons, may restore vision. The success of these approaches is dependent on the capacity of surviving circuits in the early stages of the visual system to generate and propagate an appropriate visual code in the face of neuroanatomical remodelling. To determine the capacity of surviving circuits in advanced retinal degeneration to present an appropriate visual code, we generated a transgenic mouse expressing the optogenetic actuator *ReaChR* in ON bipolar cells (second order neurons in the visual projection). After crossing this with the *rd1* model of photoreceptor degeneration, we compared *ReaChR* derived responses with photoreceptor-driven responses in wildtype (WT) mice in retinal ganglion cells and visual thalamus. The *ReaChR*-driven responses in *rd1* animals showed low photosensitivity, but in other respects generated a visual code that was very similar to WT. Furthermore, *ReaChR rd1* units in the retina had high response reproducibility and showed sensitivity normalisation to code contrast stably across different background intensities. At the single unit level, *ReaChR*-derived responses exhibited broadly similar variation in light response polarity, contrast sensitivity and temporal frequency tuning as WT. Units from WT and *ReaChR rd1* mice clustered together when subjected to unsupervised community detection based on stimulus-response properties. Our data reveal an impressive ability for surviving circuitry to recreate a rich visual code following advanced retinal degeneration and are promising for regenerative medicine in the central nervous system.

## Introduction

The ability of the central nervous system to convey and process information relies upon the properties of individual neurons and their arrangement into functional circuits. The tendency for both properties to be altered in degenerative conditions represents a fundamental challenge for regenerative medicine. Thus, as an increasing number of approaches to restore function at the primary site of lesion become available, an important question is the extent to which downstream neural circuits retain the ability to make effective use of reactivated inputs (Barker et al., 2018). Vision represents an excellent model system in which to approach this question. Firstly, the ability of the retina and brain to perform signal computations to produce a diverse and complex representation of the visual environment (the visual code) can be assessed by recording responses to carefully designed visual stimuli. Secondly, animal models of photoreceptor degeneration, such as the *Pde6b*^*rd1*^ (referred to here as *rd1*) mouse, provide well-defined defects in the input signal for vision (Pittler & Baehr, 1991; reviewed in Chang et al., 2002; Veleri et al., 2015), and these have been associated with secondary degeneration and reorganisation of retinal circuits (reviewed in Jones et al., 2005, 2016; Marc et al., 2007; Pfeiffer et al., 2020). Finally, pre-clinical methods of vision restoration are well established even in advanced degeneration (reviewed in Lindner et al., 2022; Baker & Flannery, 2018; Cehajic-Kapetanovic et al., 2022). Here, we set out to exploit these advantages to determine the capacity of visual circuits in mice with advanced retinal degeneration to generate an intact visual code.

Our approach was to express the optogenetic actuator *ReaChR* (Lin et al., 2013) in ON bipolar cells. These second order neurons are the start of the surviving visual projection in retinally degenerate mice, providing the opportunity to probe the functional capacity of surviving retinal circuitry in the absence of rod and cone photoreceptors. There is now an extensive literature showing that directly photosensitising ON bipolar cells using optogenetics is an effective approach to restoring visual responses at both electrophysiological and behavioural levels in retinally degenerate mice (Cehajic-Kapetanovic et al., 2015; Cronin et al., 2014; Doroudchi et al., 2011; Gaub et al., 2015; Lagali et al., 2008; Macé et al., 2015; Wyk et al., 2015). What remains unknown is the extent to which these approaches can recreate the visual code as generated in visually healthy animals with intact retinas. Following optogenetic targeting of bipolar cells in *rd1* mice, downstream neurons can respond to visual stimulation rapidly and resolve differences in brightness (albeit at higher light levels than required by wildtype mice). These are encouraging outcomes, but do not address perhaps the most fundamental feature of visual circuits – the ability to parse the visual scene into parallel information pathways with different feature selectivity.

The precise number of retinal information channels and what they encode varies depending on the method of classification (Coombs et al., 2006; Farrow & Masland, 2011; Sümbül et al., 2014). Baden et al. (2016) used two-photon calcium imaging to record responses of retinal ganglion cells (RGCs) to a standardised set of visual stimuli, designed to test key visual characteristics – including response polarity, contrast sensitivity, temporal frequency tuning, colour opponency, direction/orientation selectivity and receptive field size. Unsupervised clustering of these functional responses, combined with details of cell morphology, revealed at least 32 functional output channels in the mouse retina. This diversity is preserved in the dorsal lateral geniculate nucleus (dLGN), a key retino-recipient brain area in transmitting visual information to the primary visual cortex (V1), where responses reflect one of a few dominant RGC types or heterogeneous combinations of multiple RGCs (Liang et al., 2018; Román Rosón et al., 2019; Rompani et al., 2017).

The capacity of retinas with late-stage degeneration, following profound loss of rod and cone photoreceptors (Pfeiffer et al., 2020), to support a rich visual code remains largely unexplored. Some key aspects of response diversity in the fully-degenerate *rd1* retina have been indirectly assessed during studies of vision restoration, including response polarity and transience. Following optogenetic photosensitisation of ON bipolar cells, some studies find restoration of both ON and OFF responses (Cronin et al., 2014; Gaub et al., 2015; Macé et al., 2015; Wyk et al., 2015) in *rd1* animals, while others report an absence of OFF responses (Doroudchi et al., 2011; Lagali et al., 2008; Lindner et al., 2021). There is more reliable reports of variation in the degree to which responses to a light step are sustained (Lindner et al., 2021). However, a challenge in interpreting these observations is that variation in the method of transgene delivery (transduction with a recombinant viral vector) will provide different levels of expression across the target cell population. This likely imposes a potential stochastic variation in the response properties of downstream neurons.

We overcome this problem by generating a transgenic *rd1* mouse to achieve more reproducible and uniform expression of *ReaChR* across the ON bipolar cell population. We then present a battery of visual stimuli to allow comparison of visual response properties with those observed in the wildtype retina driven by rod and cone photoreceptors. Recording at the retina and dLGN, we find a remarkable concordance between the two groups. This extends to fundamental features such as contrast sensitivity, temporal frequency tuning and response reproducibility, to overall complexity of the visual code. Our work provides hope that the capacity of neural circuits to appropriately process and transmit information can be retained even following advanced degeneration.

## Results

### A transgenic model for optogenetic interrogation of retinal circuits in rd1 mice

The lack of photoreceptor input poses a challenge when assessing the ability of the remaining circuitry to process and transmit visual information in late-stage retinal degeneration. To address this, we developed a transgenic mouse model, lox-stop-lox *ReaChR-mCitrine; Grm6*^*cre*^; *Pde6b*^*rd1/rd1*^ (referred to as *ReaChR rd1* throughout), designed to express the long-wavelength-activatable variant of channelrhodopsin, *ReaChR* (Lin et al., 2013) in rod and cone driven ON bipolar cells of a degenerate retina. *ReaChR* produces highly reproducible responses with excellent temporal kinetics (Lin et al., 2013; Sengupta et al., 2016), allowing us to explore the true functional limits of the *rd1* retina. We targeted expression in ON bipolar cells as these are the earliest neurons in the surviving retinal pathway, allowing us to probe the circuitry upstream of retinal ganglion cells. We confirmed that the *Grm6*^*cre*^ driver line (Morgans et al., 2010) results in *ReaChR* expression in ON bipolar cells by co-localisation of the *ReaChR-mCitrine* fluorescent tag with staining against the rod ON bipolar cell marker, PKCalpha (Fig 1A, more detail in supplementary Fig 1).

**Figure 1.**
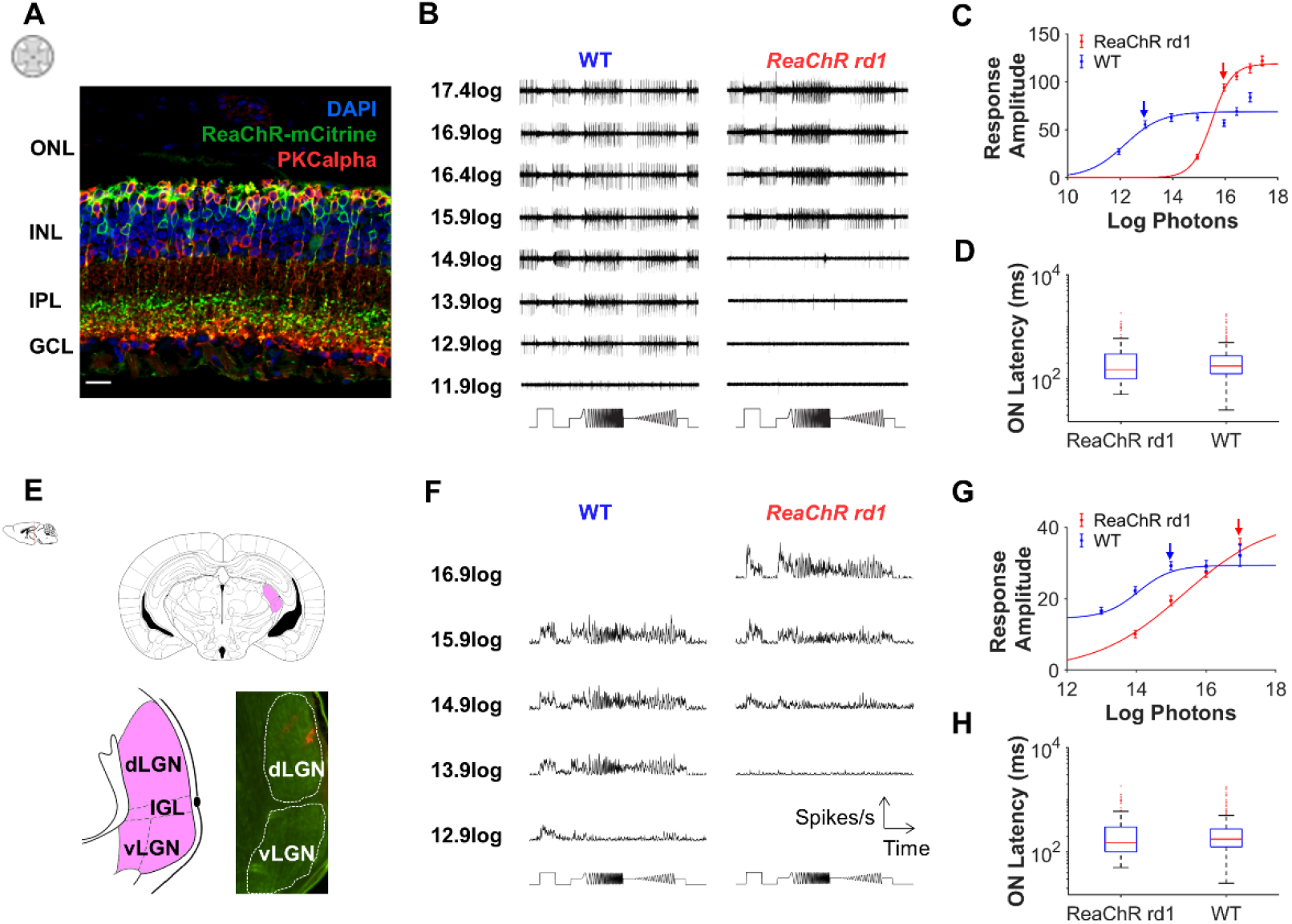
a) Retinal sections for *ReaChR rd1* mice counterstained for *ReaChR-mCitrine* tag (anti-GFP, green), rod ON bipolar cell marker, anti-PKCalpha (red), and DAPI (blue). Outer nuclear layer (ONL), inner nuclear layer (INL), inner plexiform layer (IPL) and ganglion cell layer (GCL) are labelled. Scale bar = 20µm. b) Raw data showing representative responses from multi-electrode array recordings of *ReaChR rd1* and WT retinal explants across a range of irradiances. Stimulus timing shown below. c) Irradiance response curves for transient and sustained ON units from *ReaChR rd1* (n = 414) and WT (n = 287) retinas. Irradiances used for comparison between groups are marked with arrows. d) Latency to response onset of 3s step for transient, sustained, and biphasic light responsive units (n = 480 for *ReaChR rd1*, n = 347 for WT). e) dLGN recordings. Location of LGN shown in pink on coronal section of brain (top) with details of dorsal LGN (dLGN), ventral LGN (vLGN) and intergeniculate leaflet (IGL) on bottom left. Bottom right shows histology of dLGN, with placement of electrodes in red. Brain cross-sections and dLGN diagram adapted from Paxinos & Franklin (2008). f) Representative PSTH (25ms bins) showing responses from *ReaChR rd1* and WT dLGN unit across a range of irradiances. Stimulus timing shown below. g) Irradiance response curves for transient and sustained ON units from *ReaChR rd1* (n = 225) and WT (n = 258) dLGN. Irradiances used for comparison between groups are marked with arrows. h) Latency to response onset of 3s step for transient, sustained and biphasic light responsive units (n = 261 for *ReaChR rd1*, n = 285 for WT).

### Light sensitivity, response amplitude and latency

As a first step to defining the functional capacity of visual circuits in the *rd1* mouse, we presented a stimulus comprising full-field changes in white light intensity over a range of contrasts and temporal frequencies (Fig 1B; after Baden et al., 2016) to visually intact wildtype (WT) and *ReaChR rd1* mice at 5 months of age. In separate preparations, we recorded responses at the level of retinal ganglion cells (RGCs; recordings from *ex vivo* retinal explants) or the dorsal lateral geniculate nucleus (dLGN, *in vivo* recordings performed under anaesthesia) using multi-electrode array technologies. As expected, both RGCs and dLGN neurons in WT mice responded to elements of this stimulus across a range of background light intensities (Fib 1B, C respectively). *ReaChR rd1* mice also responded to the stimulus but only at higher light levels.

To fully characterise these responses, we first quantified the change in firing rate to the simplest element of the stimulus, a full contrast 3s light step. In the retina, 51% of *ReaChR rd1* and 34% of WT units were defined as light responsive at the brightest setting tested for WT (16.95log photons/cm^2^/s). *ReaChR rd1* retinal units showed larger amplitude response than WT at this bright light (maximum baseline-normalised firing rate median for *ReaChR rd1* = 83.8 spikes/s, for WT = 51.6 spikes/s, Mann Whitney U-test, U = 6.73, p < 0.001) but were also less sensitive, with responses restricted to higher light levels (EC50 for irradiance response function = 15.5log photons). To avoid issues of response saturation, detailed comparisons between genotypes were performed at light intensities resulting in a 75% sub-saturating response in each genotype (15.9log for *ReaChR rd1* and 12.9log photons for WT). Onset of responses to the light step were faster in the *ReaChR rd1* retina at these sub-saturating conditions (Fig 1D, median response latency = 0.1s for *ReaChR rd1*, 0.15s for WT, Mann Whitney U test, U = -12.5, p < 0.001).

Units showing responses to the brightest light step were also common in the dLGN (Fig 1E) of both genotypes (54% of all *ReaChR rd1* and 50% of all WT units at 16.97log effective photons/cm^2^/s; Fig 1F). As in the retina, dLGN responses driven by *ReaChR* were less sensitive than those driven by photoreceptors in WT mice (Fig 1G). Unlike the retina, the maximal response amplitude was similar for *ReaChR rd1* and WT (median = 26.7spikes/s and 25.2 spikes/s respectively, Mann Whitney U test, U = 1.10, p = 0.269), although spike firing rates were markedly lower in dLGN compared to retina. As above, we identified 75% sub-saturating intensities for *ReaChR rd1* and WT dLGN responses (16.97log and 14.99log photons respectively). As in the retina, under these conditions, *ReaChR*-driven light responses showed faster onset kinetics than WT, with reduced response latency in the dLGN (median = 0.15s, range = 0.05-1.83s for *ReaChR rd1* and median = 0.18s, range = 0.03-1.75s for WT; Mann-Whitney U-test, U = -3.33, p < 0.001, Fig 1H).

### Step response categories

The most fundamental distinctions in visual response properties are between cells excited by increases and/or decreases in light intensity (ON vs OFF responses) and responding with either sustained or transient changes in activity. A visual inspection of step responses confirmed that this response diversity was retained in *ReaChR rd1* mice at the level of both retina and dLGN (mean peri-event stimulus histograms, PSTHs, shown at sub-saturating intensity in WT and *ReaChR rd1* in Fig 2A,E). Systematic criteria (see methods) were used to classify responses of each unit according to changes in spike firing rates observed to the beginning, end, and sustained portion of the light step. Importantly, each of the response categories found in the WT retina (transient ON; sustained ON; OFF; and biphasic ON-OFF) were also present in the *ReaChR rd1* retina (Fig 2B). While the proportion of transient ON and biphasic ON-OFF responses were similar between *ReaChR rd1* and WT retina, *ReaChR rd1* retina showed more sustained ON responses and fewer OFF responses compared to WT (Fig 2B). For a more quantitative comparison of response properties we calculated indices for transience (values close to 1 indicate maintained firing across the 3s step) and ON-OFF bias (from -1 OFF only to 1 ON only) for all units (both after Farrow & Masland, 2011). Responses were marginally more sustained for *ReaChR rd1* (median = 0.36, range = 0.02-0.94) compared to *WT* (median = 0.32, range = 0.03-0.90, Fig 2C) light responsive units in the retina, although this difference was not significant (Mann-Whitney test, U = 1.26, p = 0.26). However, there was a significant difference in ON-OFF bias with *ReaChR rd1* units significantly more biased towards ON responses (median = 0.59) compared to WT (median = 0.18, Mann-Whitney test, U = 5.86, p < 0.001, Fig 2D).

**Figure 2.**
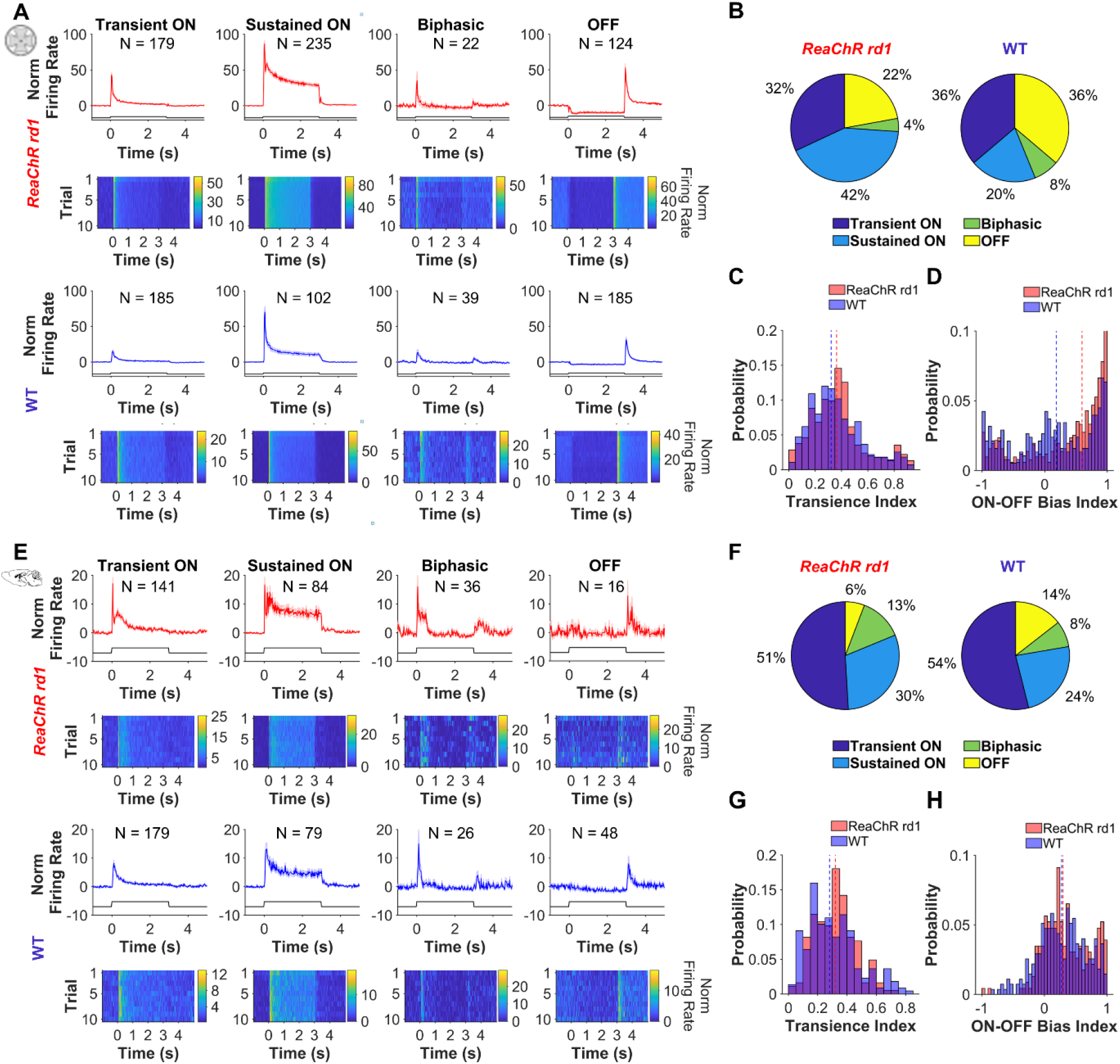
a) Mean PSTH (25ms bins, top and third row) and heat map of average response across trials (second and bottom row) in response to 3s light step (onset at 0s) for 4 different response categories detected in *ReaChR rd1* (top two rows) and WT (bottom two rows) retina. b) Pie chart showing the distribution of units across 4 different response categories in retina. c) Histogram of transience index for light responsive (LR) units in retina (n = 454 for *ReaChR rd1*, n = 275 for WT). d) Histogram of ON-OFF bias index for LR units in retina (n = 512 for *ReaChR rd1*, n = 378 for WT). e) Mean PSTH (25ms bins, top and third row) and heat map of average response across trials (second and bottom row) in response to 3s light step (onset at 0s) for 4 different response categories detected in *ReaChR rd1* (top two rows) and WT (bottom two rows) dLGN. f) Pie chart showing the distribution of units across 4 response categories in dLGN. g) Histogram of transience index for LR units in dLGN (n = 183 for *ReaChR rd1*, n = 219 for WT). h) Histogram of ON-OFF bias index for LR in dLGN (n = 230 for *ReaChR rd1*, n = 274 for WT). Dashed line on C-D and G-H shows media

At the level of the dLGN, the relative proportion of the four response categories was more comparable between *ReaChR rd1* and WT (Fig 2E), although there were notably fewer OFF units observed in the dLGN of both genotypes compared to the retina (Fig 2F). When we examined the distribution of calculated values for ON-OFF bias index, we found *ReaChR rd1* light responsive units in the dLGN were biased towards ON responses (median = 0.30 for *ReaChR rd1*, median = 0.28 for WT, Mann-Whitney U-test, U = 2.07, p = 0.039, Fig 2H), and both groups lacked the subpopulation of units with strongly OFF-biased responses seen in the retina. The transience index was comparable for WT (median = 0.32, Fig 2G) and *ReaChR rd1* (median = 0.28, Mann-Whitney U-test, U = 1.40, p = 0.163) in the dLGN.

### Contrast sensitivity and sensitivity normalisation

To assess the ability of the degenerate retina to support encoding of visual contrast, we next turned to the portion of the full-field stimulus comprising sinusoidal modulations at increasing contrast. Retinal units in both genotypes tracked this contrast chirp element at the appropriate sub-saturating intensity (representative PSTHs shown in Fig 3A, Supplementary Fig 2). Looking first at the population level response, the larger response amplitude in *ReaChR rd1* observed for the light step was recapitulated for the contrast chirp (Normalised response amplitude at highest contrast, median = 31.5 and 13.1 for *ReaChR rd1* and WT respectively, Mann-Whitney U-test, U = 8.91, p < 0.001).

**Figure 3.**
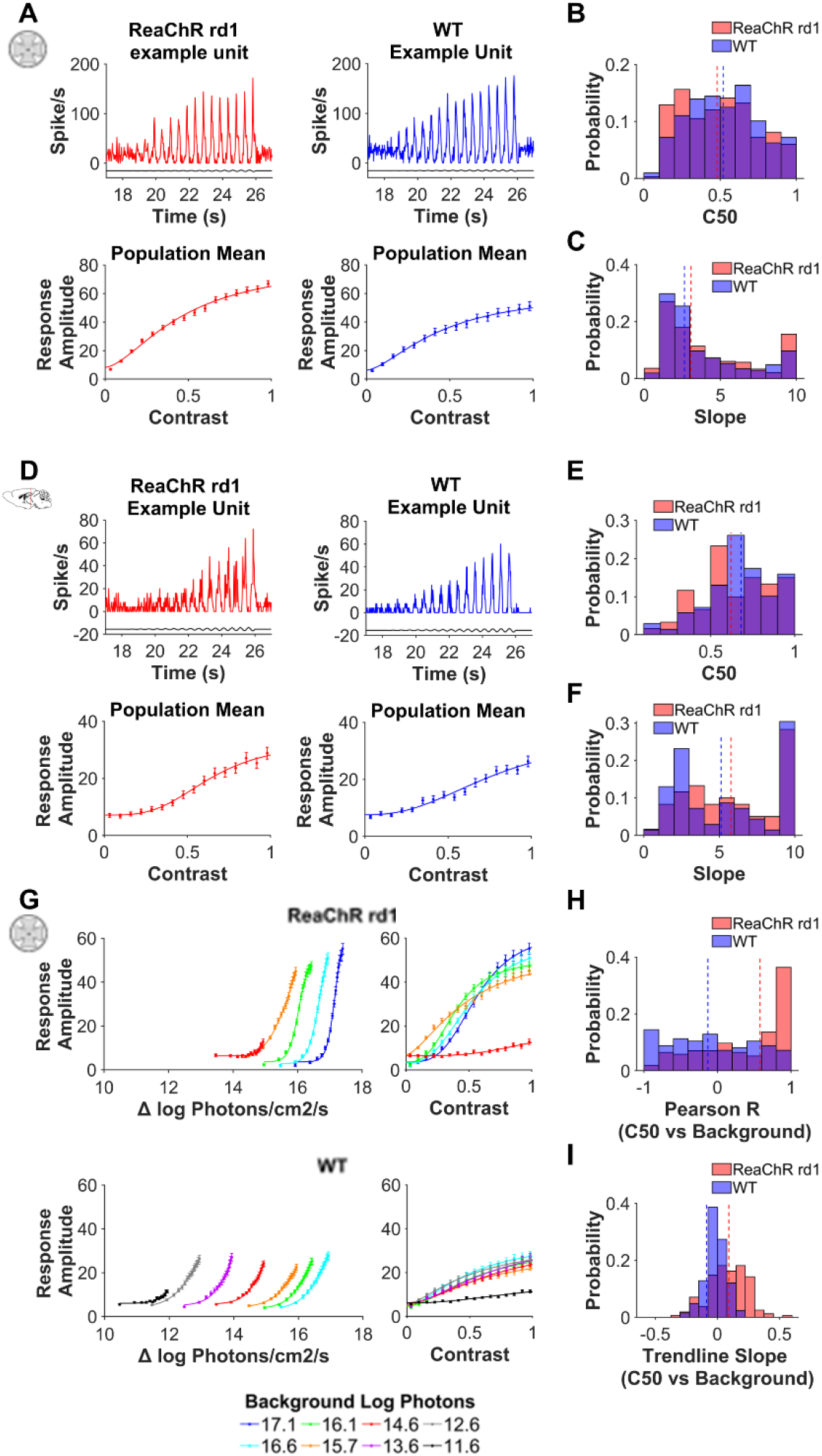
a,d) Representative example of timecourse of response to contrast chirp (Top) and contrast sensitivity function (Bottom) for individual *ReaChR rd1* (left) and WT (right) units in a) retina and d) dLGN. Timing of contrast chirp stimulus shown in black. b-c,e-f) Histogram of b,e) C50 and c,f) slope derived from best-fit contrast sensitivity function in b-c) retina (n = 333 for *ReaChR rd1* and n = 208 for WT) and e-f) dLGN (n = 60 for *ReaChR rd1* and n = 69 for WT). g) Response amplitude to contrast chirp as a function of Δ Irradiance (Left) and Michelson Contrast (Right) over different background irradiances for *ReaChR rd1* (n=303-368) and WT (n = 221-267) in retina. h-i) Histogram of h) Pearson R and i) slope of linear trendline for C50 against background irradiance for *ReaChR rd1* (n = 154) and WT (n = 124) in retina. Dashed line in B-C, E-F and H-I shows median.

More importantly, other features of the contrast-response functions across light responsive retinal units were broadly similar between the genotypes (population mean shown in Fig 3A). For both *ReaChR rd1* and WT, there was substantial diversity in the contrast required to elicit a half maximal response (C50; Figure 3B) and the slope of the contrast-response function (Figure 3C). Some units responded only to high contrast, while others showed responses across the entire contrast range (Supplementary Figure 2 for representative examples). Nevertheless, across the population there was no significant difference between genotypes in either the contrast which produced 50% response (median C50= 48% Michelson contrast for *ReaChR rd1* and 52% for WT, Mann-Whitney test, U = -1.44, p = 0.150) nor slope (median slope = 3.08 for *ReaChR rd1* and 2.64 for WT, Mann-Whitney test, U = 1.40, p = 0.162).

This finding was recapitulated in the dLGN (Fig 3D-F) where, once again, the C50 and slope of contrast response functions varied across units from both genotypes, but without a significant difference between *ReaChR rd1* and WT in either parameter (median C50 = 62% for *ReaChR rd1*, 68% for WT, Mann-Whitney U-test, U = -0.980, p = 0.327; median slope = 5.77 for *ReaChR rd1* and 5.12 for WT, Mann-Whitney U-test, U = 0.758, p = 0.449).

The ability to encode local differences in light intensity (contrast) irrespective of changes in background light intensity is a key characteristic of the visual system. To determine whether *ReaChR rd1* retinas retained this capacity, we first compared population mean contrast response relationships across background irradiances. As expected from the step response data, the *ReaChR rd1* retina responded to the contrast chirp over a narrower range of irradiances than WT (Fig 3G). Nevertheless, across this range, response amplitude could be well explained by stimulus contrast, but not by its absolute light intensity. Thus, responses to the contrast chirp at the different backgrounds were clearly separated when expressed in terms of stimulus irradiance but collapsed when expressed in terms of contrast (Fig 3G).

To quantify this behaviour and determine whether it was apparent also at the single unit level, we asked whether contrast sensitivity for each unit (as defined by C50), showed a significant relationship with background light intensity. In the case of perfect sensitivity normalisation across backgrounds, C50 should be retained irrespective of background irradiance. In practice, we found that there tended to be stronger relationship between C50 and irradiance in *ReaChR rd1* (median Pearson R = 0.58) compared to WT (median Pearson R = -0.13, Fig 3H), indicating that sensitivity normalisation was less efficient in *ReaChR rd1*. However, the slope of this relationship was relatively shallow for *ReaChR rd1* (median slope = 0.092, WT median slope = -0.088, Fig 3I), meaning that although contrast sensitivity was systematically changing with background irradiance in *ReaChR rd1* RGCs, the magnitude of this shift was relatively small at an individual level. These analyses indicate that sensitivity normalisation is occurring in the *ReaChR rd1* retina to allow effective contrast coding, albeit over a narrower range of background light intensities compared to WT.

### Temporal Frequency Tuning

We next turned our attention to the portion of the stimulus comprising an accelerating sinusoidal modulation (from 1 to 8Hz, accelerating at 1Hz/s) at full contrast. Analysis of responses to this temporal chirp component revealed that individual units in both genotypes showed strong responses to this stimulus (representative PSTHs in Figure 4A; Supplementary Fig 3). To determine, whether there was any bias in the frequencies to which the *ReaChR rd1* visual system is most responsive, we plotted normalised peak-trough amplitude as a function of stimulus frequency (shown for population mean in Fig 4A) and fit with a half-Gaussian function. In both genotypes, retinal units showed variability in preferred temporal frequency (Fig 4B). Many showed maximal response to frequencies below 2Hz, indicative of low pass tuning, but some units had larger responses to one of the higher frequencies (Supplementary Figure 3 for examples). Across the population the median preferred frequency was ∼2Hz in both *ReaChR rd1* and WT (Mann-Whitney U-test, U = 1.06, p = 0.289). Preference across this 1-8Hz range was broadly retained in the dLGN (Figure 4C-D), with median preferred frequency at ∼2Hz for both genotypes, although units in *ReaChR rd1* retina tended to be biased towards slightly lower frequencies compared to WT (Mann-Whitney U-test, U = -5.89, p < 0.001).

**Figure 4.**
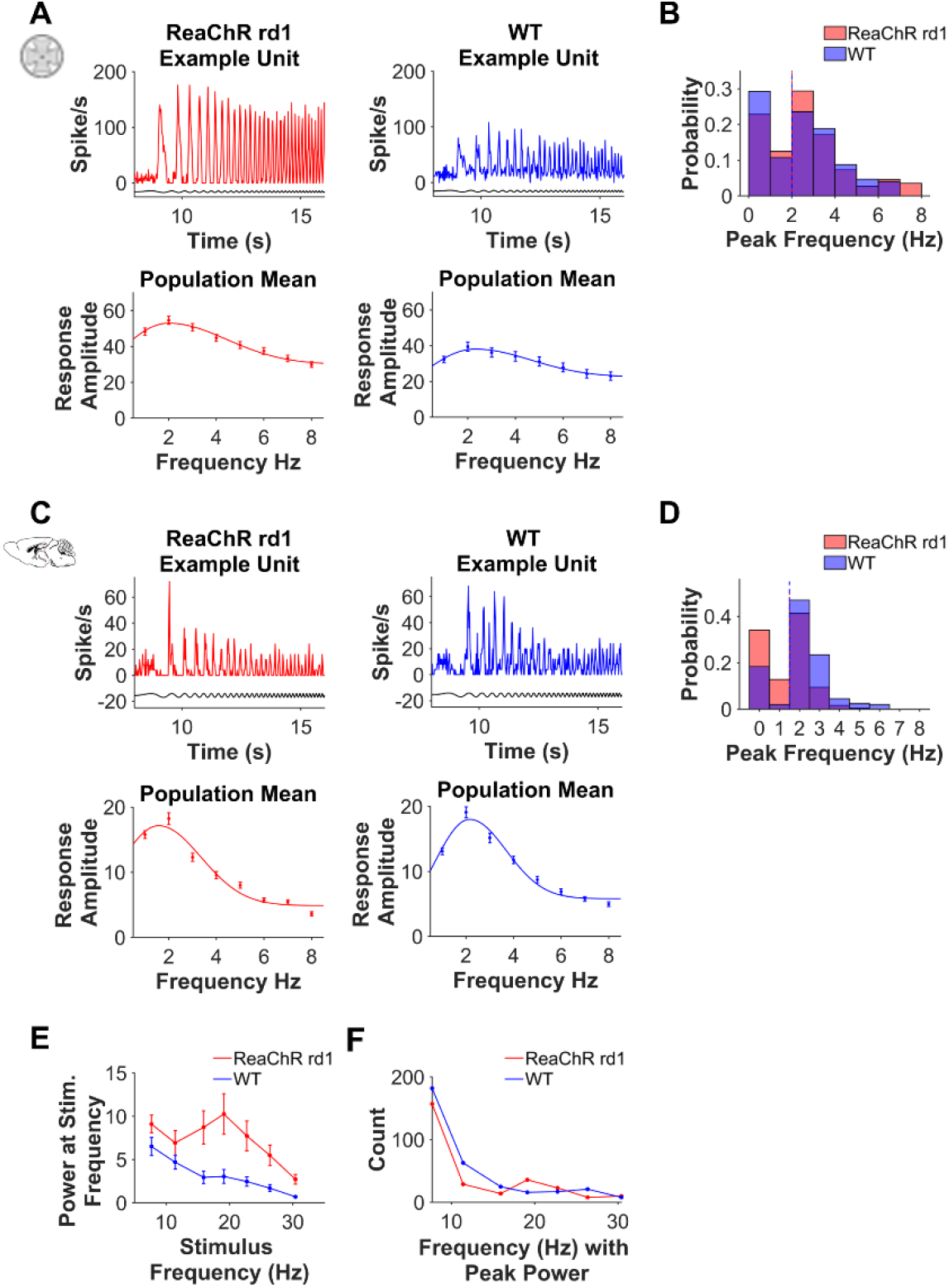
a) Representative PSTH (25ms bin size) in response to temporal chirp (Top) and temporal frequency tuning function for population mean (n = 368 for *ReaChR rd1* and n = 297 for WT, Bottom) for individual *ReaChR rd1* (left) and WT (right) retinal units. Timing of chirp stimulus shown in black. b) Histogram of peak temporal frequency (rounded to nearest integer) derived from best-fit temporal frequency tuning function (n = 368 for *ReaChR rd1* and n = 297 for WT) from retina. c) Representative PSTH (25ms bin size) in response to temporal chirp (Top) and temporal frequency tuning function for population mean (n = 179 for *ReaChR rd1* and n = 196 for WT, Bottom) for individual *ReaChR rd1* (left) and WT (right) retinal units. Timing of chirp stimulus shown in black. d) Histogram of peak temporal frequency (rounded to nearest integer) derived from best-fit temporal frequency tuning function (n = 179 for *ReaChR rd1* and n = 196 for WT). e) Mean power at different temporal frequencies (calculated from Fast Fourier transform) for all light responsive units (n = 277 for *ReaChR rd1* and n = 332 for WT) in retina. f) Distribution of units with peak power at each stimulus frequency Dashed line in B and D shows median.

In our dLGN recordings, we extended our exploration of temporal resolution to higher frequencies, presenting ∼1s blocks with full-contrast sinusoidal modulations between 8Hz and 30Hz, and calculated power for single unit firing at each stimulus frequency using Fast-Fourier transform. At a population level, for WT, the power decreased as stimulus frequency increased from 8Hz to 30Hz (Fig 4E). However, for *ReaChR rd1*, power peaked around 20Hz. When we calculated the frequency which produced peak power across this range for individual units, we found most units had weak responses to higher frequencies, with peak response at 8Hz. However, a subpopulation of *ReaChR rd1* units (59/277, 21% of units) had strong preference for higher temporal frequencies (19-23Hz, Fig 4F). Units with peak power at 19-23Hz were also present in the WT dLGN but made up a smaller proportion of the total population (33/332, 10% of units), and generally had smaller power values compared to *ReaChR rd1* (Supplementary Figure 4). Overall, the *ReaChR rd1* visual system retains the low frequency bias of WT, but with an anomalous increase in responsiveness around 20Hz in a small percentage of *ReaChR rd1* units.

### Responses to spatial patterns

Next, we examined whether there was any change in the response to spatial stimuli in the retina. We mapped spatial receptive fields in the retina using a sparse binary noise stimulus consisting of a single spot of light in a random position on a 16 × 16 grid (256 positions, Pixel area = 2.38 × 10^−3^ mm^2^, Fig 5A, as described in Lindner et al, 2021). When we compared receptive field diameters in response to sub-saturating light stimuli, we found *ReaChR*-driven responses in *rd1* retina (median = 255.3µm) were comparable to photoreceptor-driven responses in the WT retina (median = 265.8 µm, Mann-Whitney U-test, U = -1.14, p = 0.256, Fig 5B).

**Figure 5.**
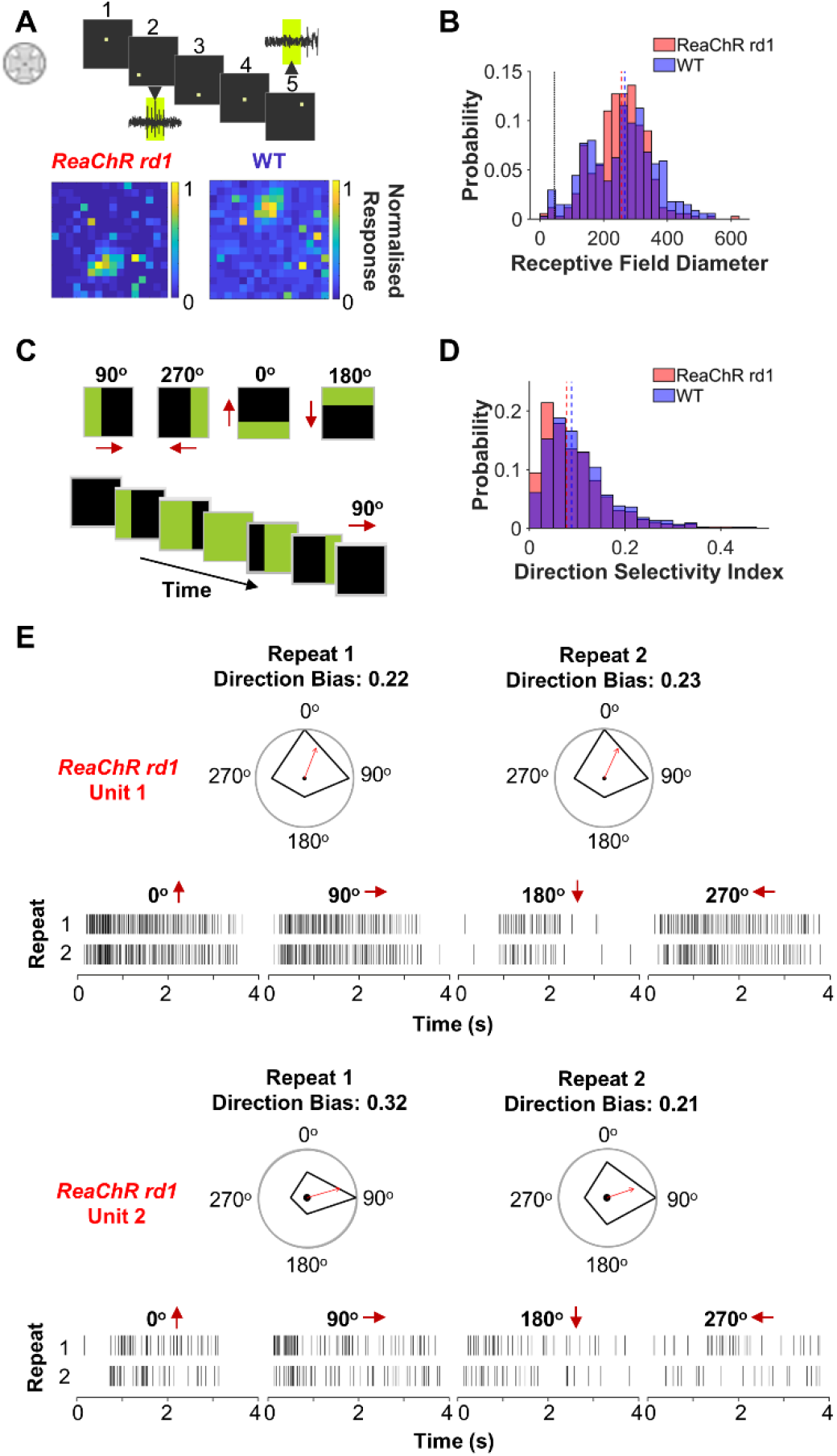
a) Receptive field maps (resolution: 2.38 × 10^−3^ mm^2^/pixel) were obtained in response to a sparse binary noise stimulus. Spike firing activity is recorded in response to different positions of the spot of light in a 16 × 16 grid (examples shown for position 2 and 5), then used to construct a receptive field map (example on right). Stimuli schematic from Lindner et al (2021). Below are example receptive fields for *ReaChR rd1* and WT retinal ganglion cells. b) Histogram of receptive field diameter (µm) for light responsive (LR) units from *ReaChR rd1* (n = 346) and WT (n = 339) retinal units. Black dotted line shows size of spot stimulus (∼42µm). c) Direction selectivity (DS) index was based on response to motion of a 565nm bar at rate of 500µm/s (total time 4s). The 565nm bar was the same width as the entire visual display. An example for motion of bar in 90° direction shown on left. This stimulus was presented for 4 directions (0°, 90°, 180°, 270°) shown on right. Red arrows show direction of moving bar. d) Histogram of DS index for LR units from *ReaChR rd1* (n = 392) and WT (n = 441) retinas. e) Raster plots for two direction selective units found in the *ReaChR rd1* retinas and polar plot of preferred direction. Two repeats are shown for each direction of moving bar, with resulting polar plot of each trial. Dashed red and blue lines in B and D show median.

We also looked at responses to moving bar stimuli (Fig 5C) to assess direction selectivity in *ReaChR rd1* and WT retinas. As expected, only a few units exhibited notable direction selectivity (Fig 5D, median direction bias = 0.08, range = 0-0.41 for *ReaChR rd1*; median = 0.09, range = 0-0.47 for WT). However, we did observe a small number also of *ReaChR rd1* cells with relatively high direction selective indices (representative examples shown in Fig 5E). Given these were relatively rare in both groups, it was not possible to determine whether direction selective cells occur more or less frequently in *ReaChR rd1* compared to WT, but their presence indicates that the capacity for direction selectivity is retained in the *ReaChR rd1* retina.

### Response reproducibility

Having defined the fundamental sensory capabilities of *ReaChR rd1*, we turned our attention to the reliability of the visual response. To this end we calculated a quality index (Baden et al, 2016) for each single unit as a measure of signal to noise across 10 trials of the full-field chirp stimulus on a scale of 0 to 1 (where 1 = identical response every trial). There was no suggestion that responses were less reproducible following retinal degeneration at either the retina or dLGN level. In the retina, in fact, the quality index tended to be higher in *ReaChR rd1* than WT (Fig 6A; median = 0.5 range = 0.10-0.96 and median = 0.27 range 0.09-0.97 for *ReaChR rd1* and WT respectively, Mann Whitney U-test, U = 9.73, p < 0.001). Overall, measures of quality index were lower in the dLGN and the difference between genotypes was no longer apparent (median = 0.17 for *ReaChR rd1* and 0.16 for WT, Mann Whitney U-test, U = 0.44, p = 0.657, Fig 6B).

**Figure 6.**
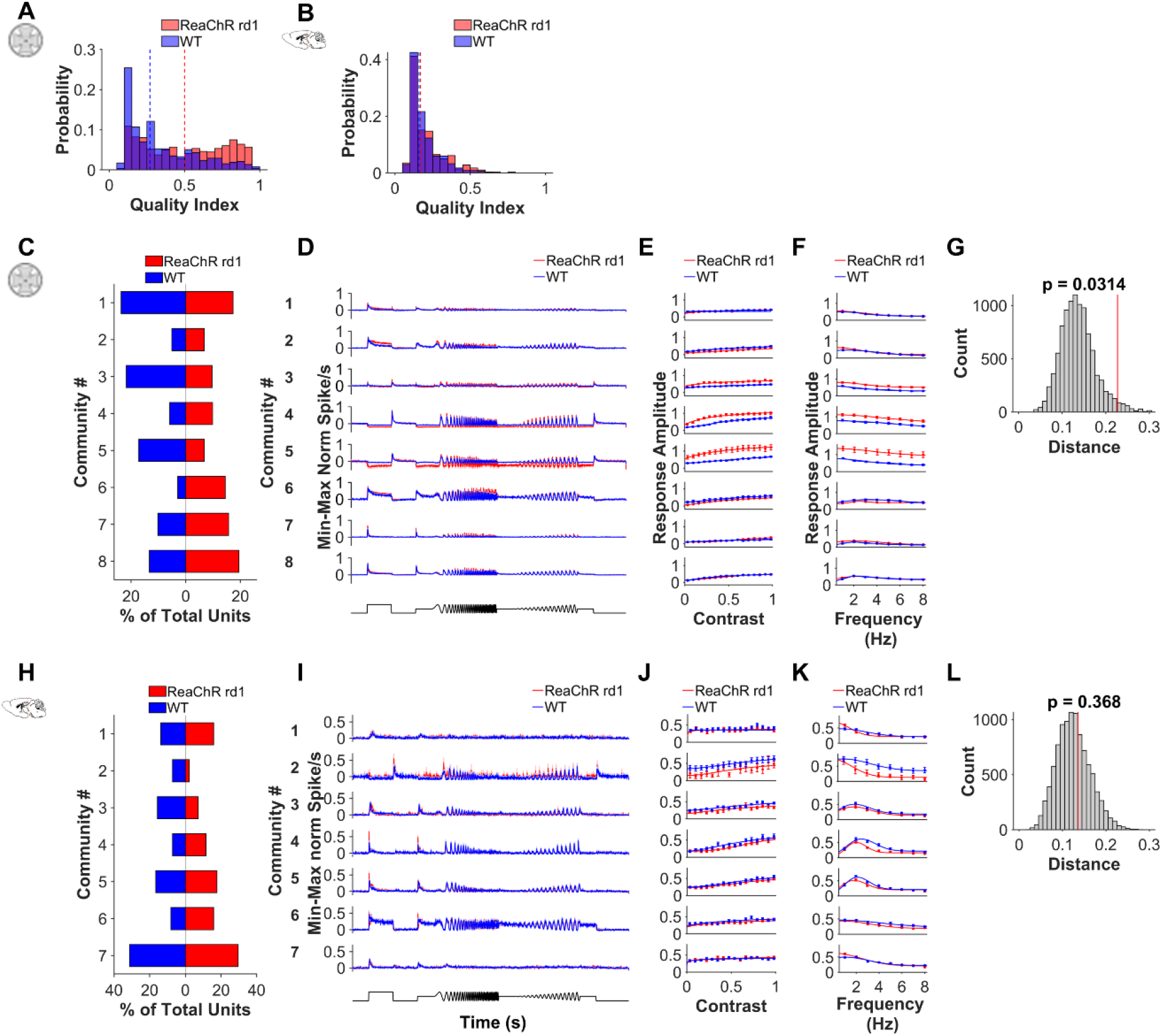
a-b) Histogram of quality index for PSTH data (100ms bins) for a) *ReaChR rd1* (n = 560) and WT (n = 511) retina units and, b) *ReaChR rd1* (n = 277) and WT (n =332) dLGN units. c,h) Distribution of units from each genotype across different communities in c) retina (n = 560 for *ReaChR rd1* and n = 511 for WT) and h) dLGN (n = 332 for *ReaChR rd1* and n = 277 for WT). d,i) Mean PSTH (25ms bins) for units with each community for each genotype in d) retina and i) dLGN e,j) Contrast sensitivity function fit to mean response to contrast chirp for e) retina and j) dLGN f,k) Temporal tuning function fit to mean response to temporal chirp for f) retina and k) dLGN g,l) Comparison of distribution of retinal units across communities. We calculated the “Distance” (Euclidian norm of the difference in the relative proportion of neurons in each community) between *ReaChR rd1* and WT (shown by red line) and compared this with a null distribution obtained by randomly shuffling replicates between two groups for g) retina and l) dLGN.

### Response Diversity and Community Detection

A key characteristic of neurons in the mammalian visual system is diversity in visual feature selectivity. Differences in the polarity and persistence of responses to the 3 sec light step (Figure 2) confirm that this diversity is retained to some extent in *ReaChR rd1*. Similarly, across the *ReaChR rd1* units in both retina and dLGN, we found considerable variability in responses to both contrast and temporal chirp elements of our full-field stimulus. We finally set out to capture the multi-dimensional nature of this response diversity for comparison between genotypes by combining probabilistic clustering and community detection.

First, response features, identified by sparse principal components analysis (Supplementary Figure 5) of pooled WT and *ReaChR rd1* responses to chirp stimulus, were grouped by using a clustering algorithm based on Gaussian Mixture Models (GMM, as in Baden et al., 2016; Caval-Holme et al., 2019). The output of this probabilistic method is referred to as clusters. Across 50 runs, the GMM algorithm returned values of between 16-24 clusters in the retina (from n = 560 WT and n = 511 *ReaChR rd1* units) and 7-11 clusters in the dLGN (from n = 277 *ReaChR rd1* units and n = 332 WT units). For most of the runs in the retina (41/50), and all runs in the dLGN (50/50), all the individual clusters identified contained units from both genotypes (Supplementary Figure 5A-B). Only 2 out of 50 runs in the retina produced a “WT only” cluster containing only WT units. This suggests that the diversity of the visual code in response to the chirp stimulus is highly preserved in both the retina and dLGN of *ReaChR rd1* animals.

However, the variability between each individual run of the probabilistic clustering algorithm prevented reliable comparison of the distribution of WT and *ReaChR rd1* units across channels and specifically the identification of functional channels that might be over or underrepresented in the *ReaChR rd1* animals compared to WT. To address this, we employed a downstream community detection analysis (Newman, 2006; Rubinov & Sporns, 2010) to obtain repeatable clustering results, while preserving the ability of the probabilistic clustering to explore different solutions. Based on a similarity matrix constructed from multiple runs of GMM clustering (up to 50), community detection was used to group units into functional output channels (referred to as communities). The community detection method reduced variability in the number of clusters and in the sets of units associated with those clusters (Supplementary Figure 6A-B, and Methods for a more detailed description of the algorithm).

Application of community detection returned 8 communities of retinal responses (Fig 6C-D). Notably, all communities contained units from both genotypes indicating that no category of response to the full field chirp stimulus was entirely absent in *ReaChR rd1*. However, there was a statistically significant difference in overall distribution of units across the 8 communities between WT and *ReaChR rd1* retinas (Fig 6G, p = 0.031, see methods for details of statistical test). Accordingly, some communities were clearly dominated by units from one genotype. For example, community 3 (p = 0.020, 67% WT units) and 5 (p = 0.004, 70% WT units) was found more often in WT animals and may be considered underrepresented in *ReaChR rd1*. Closer examination of each community (Fig 6D-F) reveals community 3 encompasses biphasic units, with transient ON and OFF responses; community 5 was made up of units with sustained ON inhibition and OFF excitation. In comparison, we found that community 6 included significantly more *ReaChR rd1* units (p = 0.001, 84% *ReaChR rd1)* with predominantly sustained ON responses. The lack of biphasic and OFF units, as well as the corresponding increase in ON units, in the *ReaChR rd1* retinas was consistent with our previous analysis of response categories (see Fig 2).

In the dLGN, 7 communities were detected (Fig 6H-K). Just as in the retina, the community encompassing OFF responses (community 2) was the most biased in favour of WT (80% WT units). However, in the dLGN there was no statistically significant difference in distribution across the categories (Figure 6L, p = 0.368). Overall, our data reveal a remarkable consistency in the categories of response to the full-field stimuli between *ReaChR rd1* and WT.

## Discussion

Regenerated light responses in *ReaChR rd1* retina and dLGN are remarkably similar to those observed in WT. Beyond the fundamental ability to reliably convey visual signals to the brain, this similarity extends to the ability to recreate much of the diversity in stimulus-response relationships that provides richness in the WT visual code. *ReaChR rd1* units show a conserved diversity of many key response types: sustained and transient; ON, OFF and biphasic; excitatory and inhibitory; high vs low contrast sensitivity; band pass vs long pass temporal frequency tuning. These characteristics are combined in such a way that when wildtype and *ReaChR rd1* units are grouped into functional output channels according to their response properties, none of the resultant groups are drawn exclusively from either genotype. We further show that spatial receptive field sizes are retained in *ReaChR rd1*, as is the ability to encode direction of movement. Finally, the ability to show equivalent responses to changes in contrast across different background irradiances reveals a capacity for sensitivity normalisation or light adaptation.

Despite the numerous similarities between *ReaChR rd1* and wildtype visual responses there were some key differences. Some of these can likely be attributed to the optogenetic actuator used to interrogate the retinal circuitry, rather than deficiencies in visual circuit performance *per se*. For example, we consistently found *ReaChR rd1* retinas only responded at the higher irradiances tested. These results undoubtably reflect the fundamentally reduced photosensitivity of *ReaChR* (Lin et al., 2013) expressed in ON bipolar cells, compared to native rod and cone photoreceptors, which have much higher photopigment content and possess signal amplifying phototransduction cascades (Arshavsky et al., 2002; Arshavsky & Burns, 2012; Fu & Yau, 2007). Similarly, the shorter response onset latency seen in *ReaChR rd1* mice may be caused by the rapid temporal kinetics of *ReaChR* (Sengupta et al., 2016), but could also be due to the cell type targeted. By expressing *ReaChR* in second order neurons (bipolar cells), the retinal circuit in *ReaChR rd1* is one synapse shorter, which could also decrease the time to onset of responses measured in retina RGCs and dLGN neurons.

The target cell type is also the most parsimonious explanation for the shift in distribution of response types. In both the dLGN and retina, *ReaChR rd1* responses increased the proportion of ON responses (particularly sustained ON responses based on results of community detection) and reduced the number of inhibitory/OFF and biphasic units. Light stimulation of rods and cones in the WT mouse retina leads to the activation of both rod and cone ON bipolar cells, as well as cone OFF bipolar cells (Dunn & Wong, 2014; Euler et al., 2014; Masland, 2012). However, light stimulation of the *ReaChR rd1* retina will likely result only in activation of rod and cone ON bipolar cells. While rod ON bipolar cells are known to connect to OFF pathways via AII amacrine cells (Bloomfield & Dacheux, 2001; Field et al., 2005; Grimes et al., 2018), the lack of direct activation of OFF bipolar cells in the *ReaChR rd1* retina may explain differences in the proportion of ON and OFF responses observed between *ReaChR rd1* and WT.

A few of the differences observed in *ReaChR rd1* compared to WT were harder to categorise, such as the larger response amplitude in the retina, the subpopulation of dLGN neurons with preference for high temporal frequencies and the finding that *ReaChR rd1* responses were highly reproducible, particularly in the retina. Our choice of optogenetic tool or use of the *Grm6*^*Cre*^ transgenic model, could all contribute to these findings, but subtle changes after retinal remodeling in degeneration could also be a factor. Particularly in the case of increasingly uniform responses seen in *ReaChR rd1* retina, it is not clear if this is necessarily positive or negative (Demb & Singer, 2015; Dunn et al., 2007; Rieke & Rudd, 2009). Under certain conditions, noise can play an important role in information handling in neuronal circuits (Rusakov et al., 2020) and losing this in the retina may result in a less naturalistic encoding of visual information. But equally, the improved signal to noise ratio could be advantageous from the perspective of vision restoration therapies.

Critically, many important visual functions found in WT were retained in the *ReaChR rd1* retina and dLGN. One exciting finding was that *ReaChR rd1* retinal units were capable of sensitivity normalization and could code contrast across a range of background irradiances. Although this was less efficient than in WT retina, it was clear the *ReaChR rd1* units were responding to relative changes in intensity, rather than simply absolute light intensity. In the intact retina, normalization to background irradiance arises from a combination of factors, including photoreceptor bleaching, adaptation in the phototransduction cascade and post-receptoral mechanisms (Demb & Singer, 2015; Dunn et al., 2007; Rieke & Rudd, 2009; Shapley & Enroth-Cugell, 1984). As *ReaChR* does not bleach (Zhang et al., 2011), the presence of this behaviour in the *rd1* retina suggests that sensitivity normalization in downstream circuitry is intact and is sufficient to enable contrast coding across different backgrounds.

More generally, we found that the diversity of responses was preserved after retinal degeneration. Critical visual features (such as contrast sensitivity, temporal frequency preference, response polarity and duration) were all captured by responses in the dLGN and retina of *ReaChR rd1* mice and varied across the population. Crucially, not only was variability in these features retained by *ReaChR rd1*, but the same functional output channels were found in both genotypes. Thus, both in our analysis of specific visual features and community detection, we found evidence that while the balance of different response characteristics may change in *ReaChR rd1* mice, no particular response type was absent. In terms of the extent to which the visual code in retinally degenerate mice is truly intact, we cannot say whether the full range of visual channels (Baden et al., 2016; Román Rosón et al., 2019) are present in the *rd1* retina without incorporating a wider array of input parameters. More significantly, future work will be required to determine whether the same classes of neurons exhibit the same stimulus-response characteristics in the photoreceptor-driven wildtype and optogenetic-driven degenerate visual system. Nevertheless, our results do suggest that the *rd1* retina in the late stages of degeneration possesses the capacity to regenerate a great diversity of visual properties.

These findings have interesting implications for our understanding of retinal reorganization in advanced retinal degeneration. To date, descriptions of reorganization have mostly focused on changes in cellular morphology and gene expression, suggesting bipolar cells (particularly their dendritic field) are most affected by the synaptic deafferentation from photoreceptor loss (Strettoi et al., 2003; Strettoi & Pignatelli, 2000). Alterations to the dendritic field of retinal ganglion cells have also been reported (O’Brien et al., 2014), although overall this cell type seems more functionally stable during degeneration (Margolis et al., 2008; Mazzoni et al., 2008). There is limited literature on the functional capacity of the degenerate retina, which has mostly focused on changes to photoreceptor-driven vision in early stages of retinal degeneration. While contrast sensitivity seems to be well preserved (Leinonen et al., 2022; Procyk et al., 2019), there is evidence of loss-of-function in some of the visual properties we tested, including attenuated response amplitude, onset latency, smaller receptive fields and an absence of sustained inhibitory/OFF responses (Procyk et al., 2019). Given that we have found the *rd1* retina retains the capacity to support these characteristics during optogenetic interrogation of bipolar cells, it is likely these findings are due to changes in photoreceptors or the first retinal synapse rather than changes later in the visual pathway.

Overall, our findings are encouraging for the prospects of regenerative medicine in the central nervous system in general, and optogenetic vision restoration therapies in particular. Targeting of ON bipolar cells in degenerate retina appears to have the potential to restore a complex and diverse array of visual responses, resulting in visual coding approximating that seen in normal WT animals. This implies that the functional capacity of deafferented circuits can be substantially retained. Our results are thus encouraging that, provided efficient transgene delivery to target cells in the retina can be achieved, optogenetic therapies could provide a high degree of visual function.

## Methods

### Animals

All experiments were conducted in accordance with the UK Home Office Animals (Scientific Procedures) Act 1986. Mice were housed under twelve-hour light / dark cycle with food and water available *ad libitum*. A new strain – *Grm6Cre; ReaChR; Pde6b*^*rd1*^ – was created for this study by breeding *Grm6*^*Cre/WT*^ (Morgans et al., 2010; MGI:4411993, shared by Robert Duvoisin & R Lane Brown, University of Oregon); *Pde6b*^*rd1/rd1*^ maintained at the University of Manchester, with *ReaChR-mCitrine* mice (Hooks et al., 2015; MGI: 5605725) obtained from Jax laboratories (strain 026294). Mice were bred to be homozygous for *ReaChR-mCitrine*, heterozygous for *Grm6 Cre* and homozygous for *Pde6b rd1* and are on a mixed C57Bl/6 x C3H background. These mice express *ReaChR-mCitrine* primarily in ON bipolar cells (Supplementary Figure 1). By 5 months old, *rd1* mice are in later stages of degeneration, following loss of rod photoreceptors and secondary cone degeneration. Prior to experiments, *ReaChR rd1* mice were kept in lighting conditions below threshold for activating *ReaChR* (Hooks et al., 2015; Lin et al., 2013; Sengupta et al., 2016). Genotyping was performed using Immomix mix red (Bioline) for Cre (Fwd 5’-TCAGCAGGTTGGAGACTTTC, Rev 5’- TTCACAACCTGTCAGACCAC, 800bp band) *rd1* (Fwd 5’-TGACAA TTACTCCTTTTCCCTCAGTCT, WT Rev 5’-GTAAACAGCAAGAGGCTTTATTG GGAA, *rd1* Rev 5’ – GCATTAATTCTGGGGCGCATG. WT 400bp, *rd1* 550bp band) and *ReaChR* (Fwd 5’ - CTTCCCTCGTGATCTGCAA, WT Rev 5’ – CAGGACAACGCCCACACA, ReaChR rev 5’ – GTTATGTAACGCGGAACTCCA. WT 96bp band, *ReaChR* 140bp band) or using an automated genotyping service (Transnetyx). For retina studies, recordings from wildtype C57Bl/6 mice (Envigo, n = 8 retinas) were supplemented with data from *ReaChR; Grm6*^*WT/WT*^; *Pde6b*^*rd1/WT*^ (n = 4 retinas), which are visually intact and do not express *ReaChR-mCitrine* or Cre recombinase in the retina. Wildtype C57Bl/6 mice bred at the University of Manchester were used for dLGN studies.

### Immunohistochemistry of retinal sections

Immunostaining of retina sections was performed as described previously (Gilhooley et al., 2022; Hughes et al., 2013). Visualization of *ReaChR-mCitrine* was enhanced using a chicken polyclonal anti-GFP antibody diluted 1:1000 that also recognizes *mCitrine* (GFP-1020, AVES labs). Anti-PKCa antibody diluted 1:1000 (ab32376, Abcam) was used to label rod ON-bipolar cells. Fluorescence images were collected using an inverted LSM 710 laser scanning confocal microscope (Zeiss) and Zen 2009 image acquisition software (Zeiss). Individual channels were collected sequentially. Laser lines for excitation were 405nm, 488nm and 561nm, with emissions collected between 440-480, 505-550 and 580-625nm for blue, green and red fluorescence respectively. Images were collected using a x40 objective with images collected every 1μm in the z-axis. Global enhancement of brightness and contrast was performed using ZenLite 2011 software (Zeiss).

### Retinal multi-electrode array recordings

Following enucleation, retinae were dissected under dim red light (>610 nm) and transferred to glass-bottomed MEA chambers (Multi Channel Systems, Reutlingen, Germany), with ganglion cell side facing down. MEA chambers (containing 252 electrodes, each 30µm in diameter and spaced 100 µm apart) were placed into the MEA recording device (MEA2100-256 system; Multi Channel Systems) and positioned within the light path of an inverted Olympus IX71 microscope. Retinae were perfused with Ames’ media bubbled with 95% O_2_/5% CO_2_ (pH 7.3) and maintained at 34 °C. Recorded signals were collected, amplified, and digitized at 25 kHz using MCS Experimenter Software (Multi Channel Systems). Retinae were perfused for at least 30 min in darkness before commencement of experiments.

A white LED light source with a daylight spectrum (Thor labs, SOLIS-3C) and an arbitrary waveform generator (RS components, RSDG2000X Series) were used to generate ‘chirp’ light stimuli (Baden et al, 2016). Intensity of light stimuli was controlled via motorised filter wheels containing neutral density filters (0-6 log units, Thor Labs). Sparse spatial noise and moving bar light stimuli were generated as described previously (Lindner et al, 2021). Briefly, a narrowband 565nm OptoLED light source (Cairn Research, Faversham, UK) and motorized filter wheels containing neutral density filters (ThorLabs, Newton, USA) were used in combination with a digital mirror “pattern stimulator” device (Polygon400, Mightex Systems, Toronto, Canada) to create sequences of defined light patterns. The power of all light stimuli (in microwatts/cm^2^) were measured at the sample focal plane using an in-line power meter (PM160T; ThorLabs), and units converted to photons per cm^2^/s using an irradiance toolbox (Lucas et al., 2014) from http://www.eye.ox.ac.uk/team/principal-investigators/stuart-peirson. All devices were automatically controlled and synchronized by a Digidata 1440A digital I/O board (Axon Instruments, Molecular Devices, San Jose, USA) and a PC running WinWCP software (J Dempster, Strathclyde University, Glasgow, UK).

Spike sorting of retina MEA data was performed using SpikeSorter software (Version 7.77b Nicholas Swindale, UBC). Raw data was filtered using a high pass 4-pole 500Hz Butterworth filter. Event detection was based on 4-5x median noise signal, with window width of 0.24ms. Results of automatic spike sorting were manually inspected and corrected using SpikeSorter software and Offline Sorter (Plexon) before being exported to Matlab for further analysis.

### In vivo electrophysiology

As described previously (Mouland et al., 2021), mice were anaesthetized by intraperitoneal injection of urethane (1.5 g/kg, Sigma-Aldrich) and placed in a stereotaxic frame. The surface of the skull was exposed and a small hole was drilled 2.2mm lateral and 2.2mm posterior to bregma. The pupil of the eye contralateral to the craniotomy was dilated using atropine (1% in saline, Sigma-Aldrich) and moistened using a small amount of mineral oil or Lubrithal (Dechra) to prevent corneal dryness. Multi-electrode arrays (A4×16-Poly2-5mm-23s-200-177-A64, NeuroNexus), consisting of 4 shanks (200µm apart) with 16 recording sites per shank, were used. Shanks were coated in CM-DiI (Fisher Scientific), positioned at 2.2mm lateral and 2.2mm posterior to bregma and inserted to a depth of 2.5-3mm targeting the dorsal lateral geniculate nucleus (dLGN). Successful recording from dLGN was confirmed by presence of light responsive units in response to 5s step of white light (5s interstimulus interval, 10 trials, 15log and 16log photons/cm^2^/s for WT and *ReaChR rd1* respectively). Once light responsive units were identified, mice were left for 30mins to dark adapt and allow neural activity to stabilise. Neural signals were acquired using a Recorder64 system (Plexon), and were amplified (x3000), high-pass filtered at 300Hz, digitised at 40kHz and stored continuously in a 16bit format. Multiunit activity was saved and analyzed offline using Offline Sorter (Plexon). In a few cases, once recording was complete, the recording probes were raised and moved ∼0.2mm posterior or anterior before insertion at an additional site, to sample a non-overlapping region of the dLGN. Once dLGN experiment was complete, mice were killed by cervical dislocation, brains removed and fixed in 4% paraformaldehyde. Single unit activity was isolated using an automated template-matching based algorithm Kilosort (Pachitariu et al., 2016) and exported as ‘virtual tetrodes’ (spike waveforms detected across 4 adjacent channels) to be manually checked in Offline Sorter (Plexon). Single unit isolation was confirmed as described in Mouland et al, 2021.

A CoolLED pE-4000 light source connected to a liquid light guide fitted with a diffuser (Edmund Optics) was used to deliver full-field stimuli, positioned at a distance of approximately 5mm from the eye contralateral to recording site. White light was used for all stimuli, consisting of output from 4 LEDs with peak emission at 385nm, 470nm, 550nm, 660nm. Irradiance was controlled using neutral density filters to produce a range from 12-16 log effective photons/cm^2^/s.

### Visual stimuli paradigms

For both retina and dLGN, responses were recorded to a standardised set of full-field stimuli (Baden et al, 2016). This consisted of a 3s step from dark to maximum light intensity, followed by 2s of dark, 2s at half maximum light intensity, an 8s temporal chirp (sinusoidal modulation between dark and maximum intensity at 1-8Hz accelerating at rate of 1Hz/s), 2s at half maximum light intensity, an 8s contrast chirp (sinusoidal modulation at 2Hz increasing from 3% to 97% contrast), 2s at half maximum intensity and finally, 3s of dark.

Due to difference in light sensitivity between WT and *ReaChR rd1*, a different range of intensities was tested for each group. For *in vivo* electrophysiology this was between 13.96-16.97 log photons/cm^2^/s for *ReaChR rd1*, while for WT, this was 12.99-16.97 log photons. For retinal recordings, this was 11.9-16.9 log photons cm^2^/s for WT, and for *ReaChR rd1* 14.9-17.4 log photons cm^2^/s.

For dLGN, responses were also recorded to higher frequency sinusoidal modulations between dark and maximum light intensity. From dark, we presented consecutive blocks of 1s of modulation at 7.8, 11.4, 15.9, 19.1, 22.7, 26.4 and 30.4Hz, followed by 3s of dark. These were conducted over same intensity range as full-field stimuli.

Receptive fields were mapped using a sparse binary noise stimulus, applied as a 16 × 16-chessboard pattern with a 48.8µm-wide pixel width (256 consecutive frames), with a frame duration of 1s. Light intensity at bright fields was either 14.2 (for WT) or 16.2 (for *ReaChR rd1*) log photons cm^2^/s, 565nm LED, contrast between bright and dark fields was 1:1000.

Direction selectivity was assessed using a sequential projection of moving wide bars into four distinct directions (0, 90, 180 and 270 degrees) at 200um/s. Light intensity was either 14.2 (for WT) or 16.2 (for *ReaChR rd1*) log photons cm^2^/s, 565nm LED, with a contrast of 1:1000.

### Analysis of Full Field stimuli – step response

Peri-event spike histograms (PSTH) were generated with bin size of 25ms. Error bars for PSTH and other graphs show standard error of the mean, unless otherwise specified. Light responsive units were identified using confidence limits test based on responses to the 3s step of the chirp stimulus: units were classified as significant if spike firing rate during the response window was greater than 2 standard deviations above (excitation) or below (inhibition) mean firing rate during baseline window – equivalent to 95% confidence limit. We used 3 different confidence limit (CL) tests; activity during the first 2s of the chirp step compared with 1s before step onset (transient ON), activity during 0.5s after step offset compared with last 1s of step stimulus (OFF) and activity during last 1s of step stimulus compared with 1s before step onset (sustained ON). We classified light responsive units into 4 different categories based on outcome of these confidence limits tests: 1) Transient ON units were significant for transient ON test, with no sig. excitation for sustained ON and no sig. response for OFF test, 2) Sustained ON units were sig. for transient and sustained ON tests with no sig. excitation for OFF test. 3) Biphasic units were sig. excitation for transient ON and OFF tests, with no sig. response to sustained ON test. 4) OFF/Inhibitory units had significant inhibition for transient and sustained ON tests and had significant responses with OFF test. Units with a quality index below 0.1 (Baden et al, 2016), a measure of response variability between repeated trials, were excluded from further analysis. Firing rate was then normalised to baseline by subtracting the average activity during 2s before step onset.

Irradiance response curves were constructed based on maximum normalised firing rate during 3s step by fitting Hill slope curve with 4 best-fit parameters (Top, Bottom, logEC50 and Slope) identified using non-linear regression. Irradiances which produced response closest to 75% maximum were identified and used for comparisons between WT and *ReaChR rd1* groups. The 75% threshold was used as it would provide sub-saturating responses in the majority of units, while ensuring good signal to noise ratio and as large a sample size of responsive units as possible.

Equation for Hill slope curve is

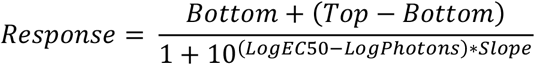

where LogPhotons is intensity in log photons/cm^2^/s

ON:OFF Bias index and Transience index were calculated using previously described methods (Farrow & Masland, 2011; Lindner et al., 2021). Latency for onset of 3s step was defined as timing of first bin where response was outside the confidence limits for the transient ON confidence limit test. Light responsive units that did not respond to onset of 3s step (ie: OFF units) were excluded from this analysis.

### Analysis of Full Field stimuli – contrast sensitivity

The response amplitude (maximum – minimum normalised firing rate during each period of the contrast chirp stimulus) was normalised to baseline activity during 1s before contrast chirp onset for each unit. This was plotted against Michaelson contrast and then fit using a contrast sensitivity function, or Naka-Rushton curve (Albrecht & Hamilton, 1982), using least-squares minimisation to identify 4 best-fit parameters (top, bottom, C50 and slope).

Equation for Naka-Ruston curve is

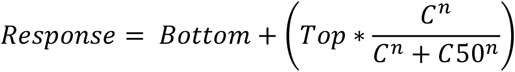

Where *n* = slope, *C* = Michelson contrast and C50 is contrast that produces half maximum response. C50 was constrained between 0 and 1, and slope was constrained between 0 and 10. Only curve fits with R^2^ > 0.5 and spiking in >10% of bins were used for comparison of contrast sensitivity parameters.

For analysis of contrast coding, the change in irradiance between peak and trough of chirp modulation for each period of contrast chirp stimulus (Δ Irradiance). Response amplitude for contrast chirp (calculated as above) at each background irradiance was then plotted against Michelson contrast or Δ Irradiance for light responsive units. Mean contrast response amplitudes were then fit with contrast sensitivity function (for amplitude vs contrast) or Hill-slope curve (for amplitude vs Δ Irradiance).

For analysis of contrast coding at an individual level, we first identified light responsive units that had contrast sensitivity functions with R^2^ > 0.5 for each background irradiance. We then calculated the correlation between C50 from contrast sensitivity curve and background irradiance, as well as the slope of the linear trendline.

### Analysis of Full Field stimuli – temporal frequency tuning

For analysis of temporal chirp, mean response amplitude (calculated as above) for each temporal frequency was fit with a half-Gaussian model (Grubb & Thompson, 2003) using least-squares minimisation to identify 5 best-fit parameters (low baseline, high baseline, Gaussian spread, peak response and peak frequency).

Equation for Half Gaussian is:

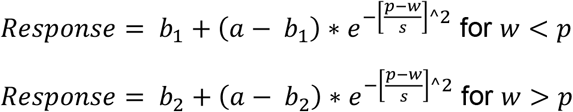

Where *w* is the temporal frequency (Hz), *p* is the temporal frequency (TF) that produces peak response, *a* is the maximum response amplitude at optimum TF, *s* is the Gaussian spread, *b1* is the baseline for frequencies lower than peak TF, *b2* is the baseline for frequencies greater than the peak TF. Only curve fits with R^2^ > 0.5, spiking in > 10% of bins and Gaussian spread > 0.51 were used for comparison of temporal frequency tuning parameters. Peak temporal frequency was rounded to nearest integer to address the limited resolution of temporal frequency analysis (sample every 1Hz).

For analysis of faster frequencies, for each stimulus frequency, we generated a PSTH with data in 10ms bins and performed a fast-Fourier transform. We then extracted the mean power for the 3 frequencies closest to stimulus frequency and plotted this as a function of stimulus frequency.

### Receptive Field mapping and assessment of Direction Selectivity

Receptive field diameter was calculated using the method described in Lindner et al. (2021) with minor modifications: For each stimulus frame, PSTHs were calculated for an interval starting 500ms before the beginning of frame, to 500ms after the end of frame (bin size 250 ms). The absolute peak difference in spike firing rate over each PSTH was then calculated, which enabled analysis of ON, OFF, ON-OFF and sustained units likewise. Resulting values were multiplied with a 2D binary matrix representing the stimulus pattern during that particular frame. Thereby, observed responses could be matched to individual stimulus locations. Resulting matrices were then collapsed and normalized. Receptive field centre positions were detected, and the width of the receptive field was determined by Gaussian fits. Receptive field centre positions were detected on smoothed data to minimize the effect of possible noise. Receptive fields which had their calculated maximum at the edge of the field of view were excluded from analysis.

The Direction Selectivity Index was calculated as described by Farrow and Masland (2011): For each responsive units, spike firing rate during the presentation of the moving bar were computed for each of the four directions of movement independently. A direction selectivity vector for each responsive unit was computed. The length of the vector was normalized resulting in the Direction Selectivity Index.

For spatial stimuli, units were identified as light responsive based on response to a 1s flash. Units were classified as light responsive if there was a change in spike firing rate of >10 spikes/s and a relative change in number of spikes > 20% during flash compared to baseline.

### Response reproducibility and diversity analysis

To assess reproducibility across trials, the quality index (Baden et al, 2016) was calculated for PSTH data in 100ms bins.

Sparse Principal components were generated for the following windows of chirp stimulus – Step (0.5-4.5s), temporal chirp (6.5 – 15.5s) and contrast chirp (17.5-24.5s) – using the SPaSM toolbox (Sjöstrand et al., 2018), as described in Caval-Holme et al (2019) and Baden et al (2016). This allows the extraction of response features that are localised in time. We pooled mean PSTH (25ms bins) for all light responsive units from both groups and extracted up to 30 features with 5 non-zero time bins. We then discarded those that accounted for < 1% of the variance. Response features that met these criteria for each window were then combined to produce a total of 21 features for retina data and 30 features for dLGN.

sPCs from pooled *ReaChR* and WT data were then clustered with a mixture of Gaussian models, a probabilistic model using random initialisation. According to this method, the optimum number of clusters is determined based on the lowest Bayesian information criteria, which rewards fit but penalises complexity, and a Bayes factor below 6, as used by Caval-Holme et al (2019), as a threshold for when there was no longer evidence for further splitting.

The probabilistic clustering was repeated 50 times. A pairwise similarity matrix was generated based on how frequently each unit pair clustered together (with 0 being never clustered together and 1 being always clustered together). Based on this similarity matrix, a community detection algorithm (Newman, 2006) using the Brain connectivity toolbox (Rubinov & Sporns, 2010 at https://sites.google.com/site/bctnet/) was used to group units into functional output channels.

To compare distribution of units across communities between two groups, we calculated the distance (Euclidian norm of the difference in the mean relative proportion of neurons in each cluster) between *ReaChR rd1* and WT (shown by red line in Fig 6G and 6L) and compared this with a null distribution obtained by randomly shuffling retinal recordings between two groups 10,000 times. To compare proportion of units from each group within each cluster, we first calculated the % of total units from each group in a given community for each recording, and we used these averages to calculate the percentage difference between *ReaChR rd1* and WT. This percentage difference was compared with a null distribution generated as above.

---

**Supplementary Figure 1.**
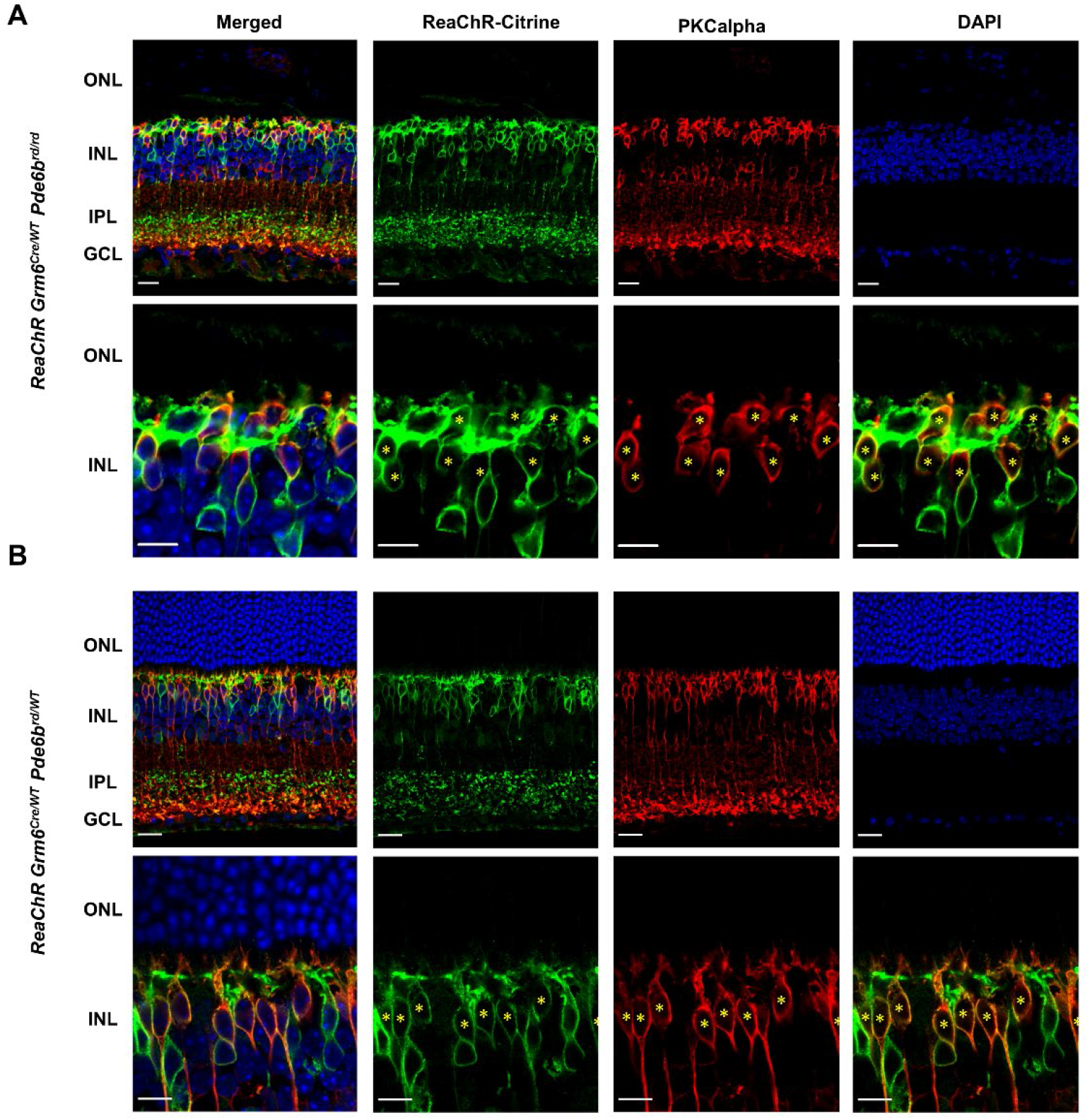
a-b) Retinal sections for a) retinally degenerate *ReaChR Grm6Cre rd homozygous* and b) non degenerate *ReaChR Grm6Cre* rd heterozygous mice counterstained for *ReaChR-mCitrine* tag (anti-GFP, green), rod-ON bipolar cell marker, anti-PKCalpha (red), and DAPI (blue). Scale bar = 20µm for top row, 10µm for bottom row. * indicates PKCalpha labelled rod ON bipolar cells also labelled for *ReaChR*.*mCitrine*. Note that *ReaChR mCitrine* expression is also detected in non-PKCalpha labelled cells consistent with expression in cone ON bipolar cells (Euler et al., 2014).

**Supplementary Figure 2.**
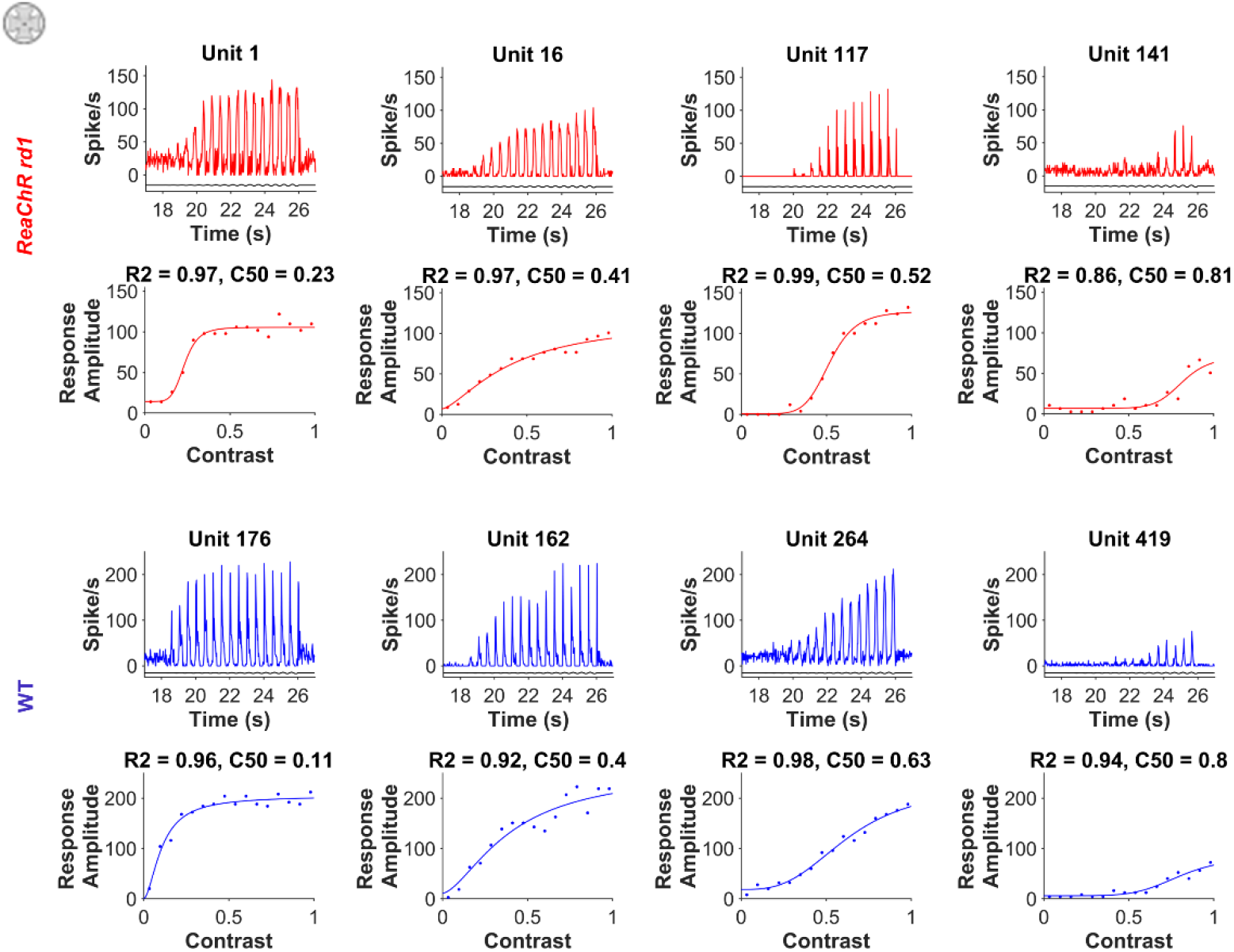
PSTH (top) and contrast sensitivity function (bottom) for 4 example units from a) *ReaChR rd1* (red) and b) WT (blue) retinas, showing diversity in C50 and slope within each genotype.

**Supplementary Figure 3.**
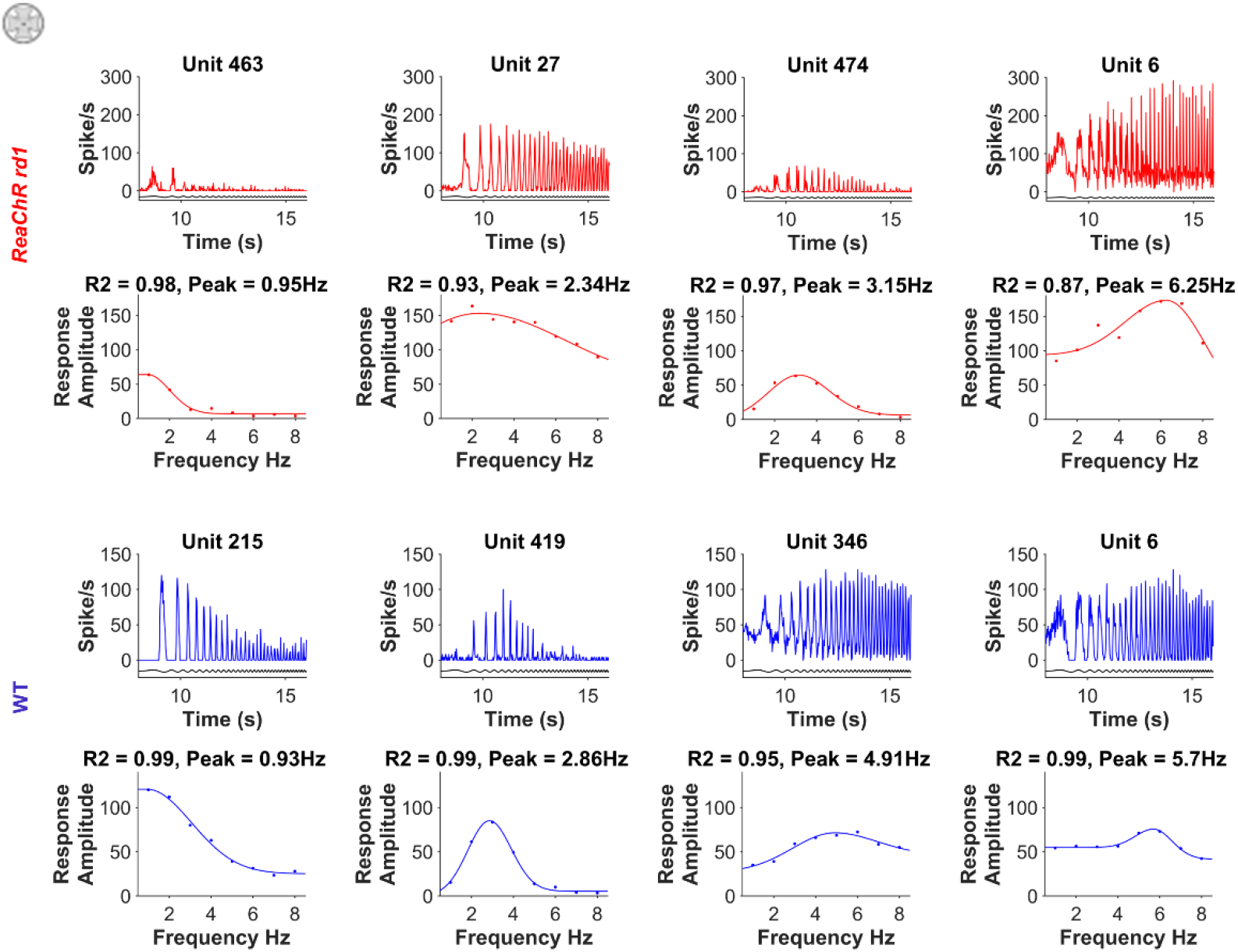
Time course (top) and temporal frequency tuning function (bottom row) for 4 example units from a) *ReaChR rd1* (red) and b) WT (blue) retinas, showing diversity in peak frequency and bandwidth of tuning curves within each genotype.

**Supplementary Figure 4.**
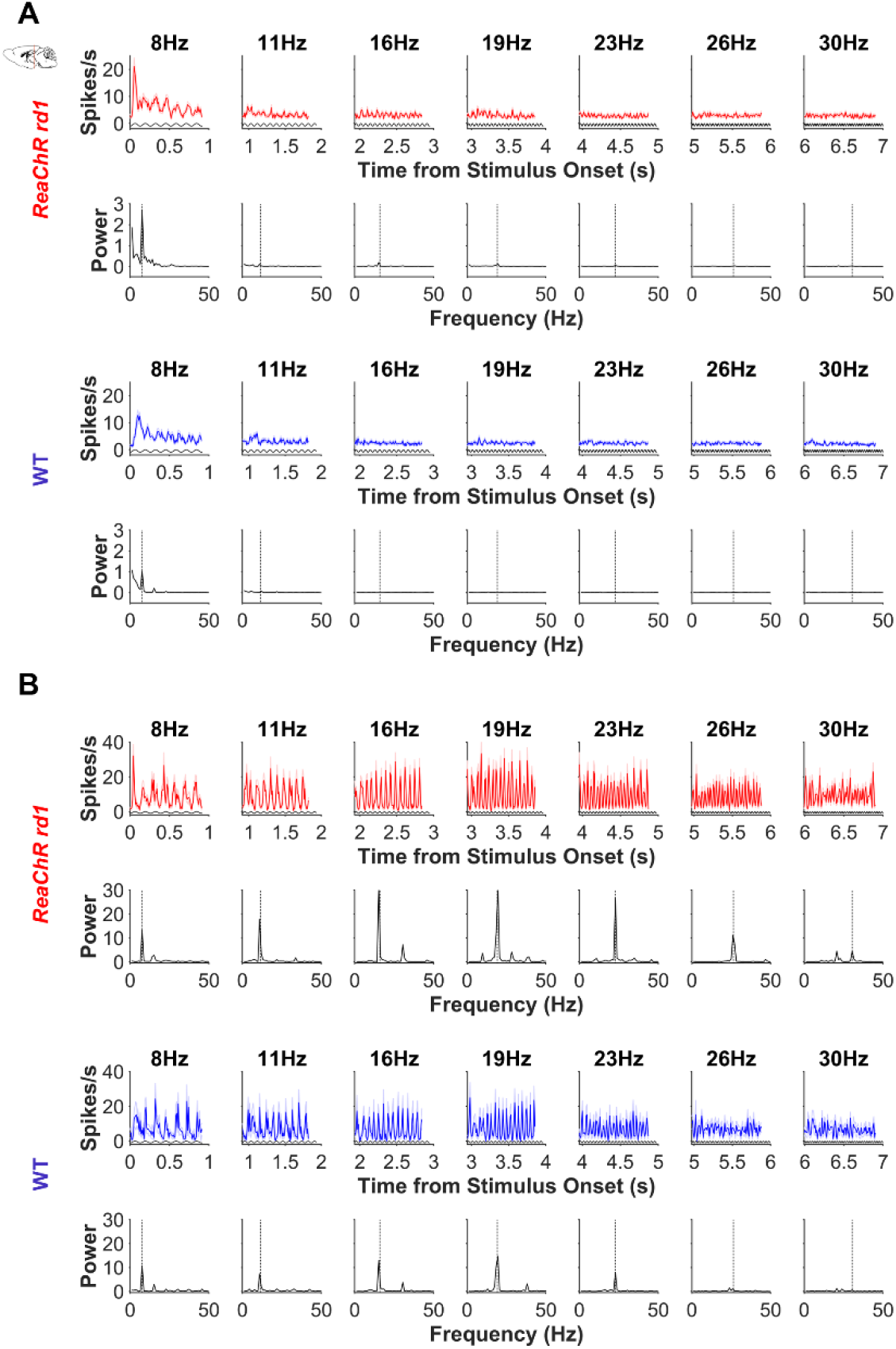
Mean PSTH (top, 10ms bins) and periodogram (bottom) at each stimulus frequency (labelled above PSTH) for all dLGN units with peak power at either a) 8Hz (N = 157 for *ReaChR rd1* in red, N = 182 for WT in blue) or b) 19Hz (N = 36 for *ReaChR rd1* in red, N = 16 for WT in blue). Stimulus timing shown in black on PSTH. Dotted vertical line on periodogram shows stimulus frequency.

**Supplementary Figure 5.**
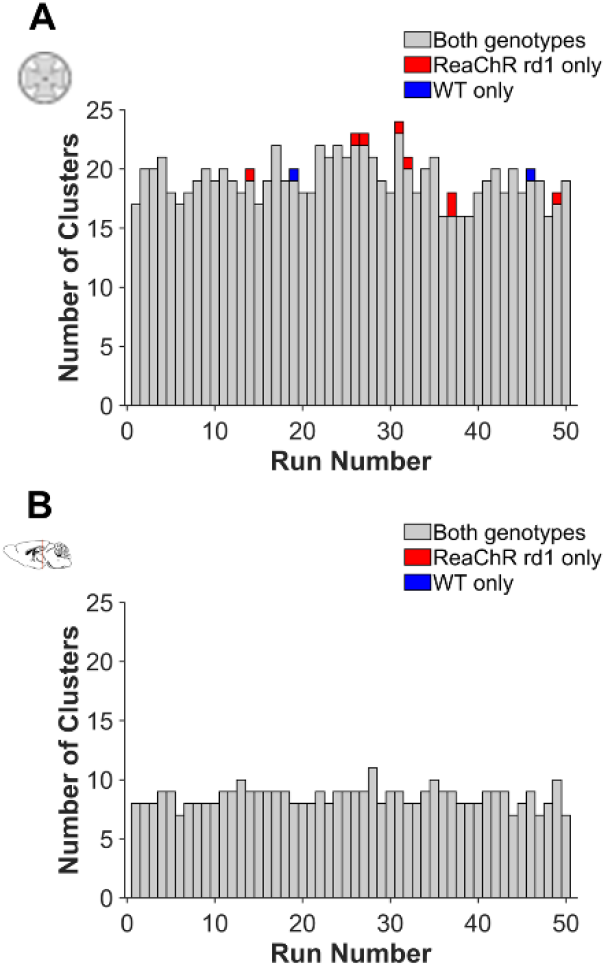
Summary of 50 runs of clustering based on Gaussian Mixture Models (Baden et al., 2016). Bar heights show the total number of clusters generated for each repeat of clustering algorithm, with different colours indicating the relative number of clusters that contain units from both genotypes (grey), *ReaChR rd1* units only (red) or WT units only (blue). Shown for a) retina and b) dLGN data.

**Supplementary Figure 6.**
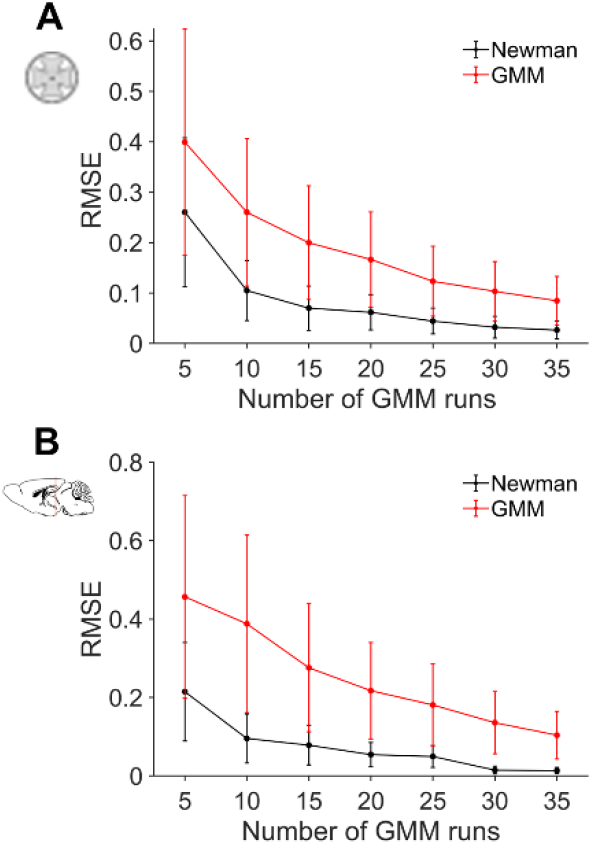
Community detection increases the reliability of Gaussian Mixture Models clustering. a-b) Root mean square error (RMSE, Mean±SD) across similarity matrices obtained by sampling an increasing number of repeats of the GMM clustering algorithm (red lines). RMSE decreases as function of the number of GMM repeats sampled, reported on the x-axis. At each repeat of the clustering algorithm, the associated similarity matrix was also employed to detect communities by using the Newman community detection method (Newman, 2006; Rubinov & Sporns, 2010), and a new similarity matrix was generated based on such communities. RMSE (Mean±SD) was then recalculated based on the Newman-derived similarity matrices (black lines). Note that the application of community detection (black lines) substantially reduces the RMSE. Shown for a) retina and b) dLGN data.

